# Nationwide survey on the barriers to converting turfgrass lawns to pollinator-friendly native wildflowers

**DOI:** 10.1101/2020.06.02.129452

**Authors:** Nash E. Turley, Joshua Hogan, Gloria J. Diehl, Aaron C. Stack, Barbara J. Sharanowski

## Abstract

The abundance and diversity of insect pollinators around the world is declining and habitat loss is a leading cause. Turfgrass lawns cover a vast area in North America and provide a great opportunity for habitat restoration to native wildflowers by the general public. Efforts to encourage the public to replace lawns with wildflowers could be improved by a better understanding of the thoughts and opinions of the public about lawns. We conducted a nationwide online survey to understand what barriers are most important in preventing people from converting a 6 x 6 ft portion of turfgrass lawn to native wildflowers. We also collected data on a variety of demographic factors to see if those influence survey responses. Over 3200 people took survey across the US. We found that ‘Maintenance time’ and ‘Not knowing what to do’ were the most important barriers to creating wildflower habitat. Age was the most important demographic factor impacting results with young people significantly more likely to select multiple barriers in the survey. For example, people aged 18-34 were 4.3 times more likely to indicate ‘Maintenance cost’ would prevent them from creating a wildflower plot than those age 65 or older. Those who had already created a wildflower plot, or those who were members in a native plant or pollinator organization were less likely to select barriers across the board, except for external barriers related to homeowners associations, neighbors, and local governments. This shows that these are persistent concerns even for those that are otherwise keen to create wildflower habitat. Our results suggest that outreach promoting pollinator-friendly native plant gardens should focus on clear and simple methods, small plots that will not take too much time and less likely to provoke neighbors or authority figures.

## Introduction

Human activities on earth have resulted in a loss of biodiversity worldwide [1, 2]. For example, there has been a steady decline in the abundance of birds in North America over the last 50 years with habitat loss thought to be the single most important cause [3]. The abundance and diversity of native insect pollinators, such as bees, flies, and butterflies are also falling [4–7]. This is particularly troubling because these pollinators provide significant ecological and economic services, and their decline would result in serious negative impacts worldwide [8–10]. The leading cause for pollinator declines is also habitat loss, notably the loss of native wildflowers [4, 9] and the overuse of pesticides and fertilizers [6, 7]. Turfgrass lawns contribute to this problem as they provide little to no resources for pollinators [11] and cover a total surface area of 163,812 km^2^ in the US, roughly the size of Georgia [12]. This is three times larger than the area of irrigated corn, the largest irrigated crop in the US [13, 14]. Turfgrass is typically grown as a monoculture and has several negative environmental impacts associated with maintenance and upkeep. A ‘pretty’ lawn often requires large chemical inputs such as fertilizers, insecticides, and herbicides, and extreme water use which accounts for up to 75% of a household’s total water consumption in semiarid regions [12].

While turfgrass lawns have become the standard for home landscapes, they have great potential to be transformed into landscapes rich with native wildflowers that are beautiful, sustainable, and better for pollinators [15, 16]. However, the widespread adoption of this type of habitat restoration by private landowners pose several cultural challenges. As there are strong cultural norms associated with mowed turfgrass lawns in residential areas [17–19], turfgrass is well-liked by many homeowners and for some represent neatness [20], wealth, and security [21, 22]. That said, yards with mixtures of turfgrass and gardens of native plants have been found to be equally as attractive as traditional turfgrass lawns [23]. Therefore, outreach efforts could be successful at convincing the average homeowner to transform a portion of their turfgrass lawn into a garden of native wildflowers, but these efforts need to be guided by an understanding of the factors that influence people’s decisions.

Larson et al (2016) surveyed homeowners about what they valued in their lawns and found that low maintenance and aesthetically pleasing designs were preferred [18]. Dahmus and Nelson (2014) conducted a similar survey asking homeowners how they conceived their yard to be part of the local ecosystem [24]. They found that the surveyed individuals do have a complex understanding of their yard as part of the local ecosystem, but that they usually had some prominent gaps in their understanding, most notably knowledge of biodiversity and ecosystem services. Here we build upon this body of research by conducting a nationwide survey to better understand the general public’s thoughts regarding converting portions of their turfgrass lawns to wildflowers. More specifically, we sought to answer two questions:

1. What barriers are most important in preventing people from converting patches of lawns to wildflowers?
2. How do barriers that prevent lawn to wildflower conversions vary by demographic factors and personal characteristics?

We chose to target this survey towards an audience that is already interested in plants and pollinators and likely concerned about pollinator declines. We focused on this audience for three reasons. First, they ended up being the people that were most likely to take the survey. Second, they represent the people that are most likely to act and create wildflower habitat. Third, we wanted the results of the survey to help guide our outreach efforts for our public science project Lawn to Wildflowers (https://lawntowildflowers.org) and people that are already interested in wildflowers and pollinators are who we will reach out to first. This project provides resources to help the general public convert lawns to pollinator-friendly wildflower habitats, to learn to identify pollinators, and to collect data on pollinators.

## Methods

### Survey details

We developed an online survey using the Qualtrics^XM®^ Software (https://www.qualtrics.com/) that was distributed between 28 August, 2018 and 18 April, 2019. Anyone could take the survey, though we requested that the individuals only take it if over the age of 18. The survey included a question which asked people to select all the barriers that might prevent them from converting a 6 x 6 ft patch of turfgrass in their front yard to a patch of wildflowers. We included 11 potential barriers to lawn-to-wildflowers restorations which we chose after talking with colleagues promoting native plants and pollinator-friendly landscaping, and surveying related literature [18]. These barriers can be placed in three categories: individual barriers based on personal opinions and circumstance (‘Appearance’, ‘Maintenance cost’, ‘Maintenance time’, ‘Loss of space for recreation’, and ‘Not knowing what to do’), external barriers relating to plants and animals (‘Undesirable plants’, ‘Undesirable wildlife’, and ‘Bee stings’), and external barriers relating to other people (‘Fines or infractions from local government’, ‘Opinions of neighbors’, and ‘Violation of homeowners association policies’). We also included a twelfth choice as ‘None apply.’

The survey also included a series of multiple-choice questions relating to the demographics of the respondent. This included basic demographic questions related to age, gender, income and education. We pulled nationwide census data from 2018 to compare the demographics of our respondents to the nation at large (S1 Table). We included some additional questions related to the topic of lawns, homeowners’ associations, membership in plant and pollinators groups, and if they had already created a wildflower plot. Finally, we also asked two additional Likert Scale questions to assess their likelihood of creating a wildflower plot and to identify their concerns about pollinator declines. The full text of the survey is available in supplementary information (S1 Appendix). The Qualtrics software determined the approximate Latitude and Longitude for most of the survey responses which we used to create a map of where responses came from (S1 Fig.). We included a question that asked for Zip Code, using the middle of the Zip Code area to determine location when other coordinates were not available. Responses that were incomplete, did not have location data, or were from outside the US and southern Canada (below 54° N) were removed from the dataset (N=483).

### Distribution of survey

We promoted the survey using our Lawn to Wildflowers social media accounts, utilizing paid advertisements and boosted posts on Facebook and Instagram. A sample of the post we used for most paid advertisements is included in supporting information (S2 Fig.). One round of advertisements targeted people in the US by using the topic keywords “pollination, beekeeping, wildflower, and lawn” to reach individuals already interested in the topics of the survey. To diversify the audience taking the survey we also targeted a younger audience (below 50) and more conservative audience using the topic keywords “American football, lawn mower, and lawn”. We also conducted an email campaign where we messaged Native Plant Society chapters in every US state or region and encouraged them to share the survey with their members.

### Statistical analyses

To determine the most important barriers that may prevent lawn to wildflower conversions, we simply compared the differences in counts in responses to each of the 12 possible choices. For visualization, we converted counts to percentages of total respondents. To test if demographic factors and personal characteristics influenced those results, we conducted Chi-Squared tests for independence. We looked at age, gender, income, education level, membership in a homeowner’s association, membership in a native plant or pollinator group, and if they had already created a wildflower plot. Because we conducted a large number of tests, which increased the possibility for Type 1 error, we chose to focus on results with P<.001. We excluded the following demographic categories that had too few respondents (< 40) as we felt the small sample size would not be a reliable representation of the group: when looking at gender we excluded gender nonconforming, when looking at income we excluded those who made >$500,000 per year, and when looking at education level we excluded those who did not complete high school. The full dataset used in the analyses is available in suporting information (S2 Appendix).

## Results

Our final dataset had 3249 survey responses located across the US and some in Southern Canada. Most survey responses originated from the eastern US, most notably the coasts of Florida and New England, although responses were scattered throughout the US (S1 Fig.). Our surveyed population tended to be older, more educated, and more female than the average person according to US census data. Of our respondents, 56.5% were over the age of 55 (Table 1), compared to only 28.9% of US citizens in the same age range (S1 Table). Also, 80.8% of our respondents had achieved some degree of formal college education (defined as an associate degree or higher) as opposed to 41.2% of Americans in census data (S1 Table). Furthermore, 76.7% of respondents identified as women, while 50.8% of Americans overall identify as women (Table 1, S1 Table). Finally, our audience had a very strong interest in the topic, with 71% of respondents identifying that they were extremely concerned about pollinator declines and 79% identifying that they would create a 6 x 6 ft wildflower plot (Table 2).

**Table 1.**
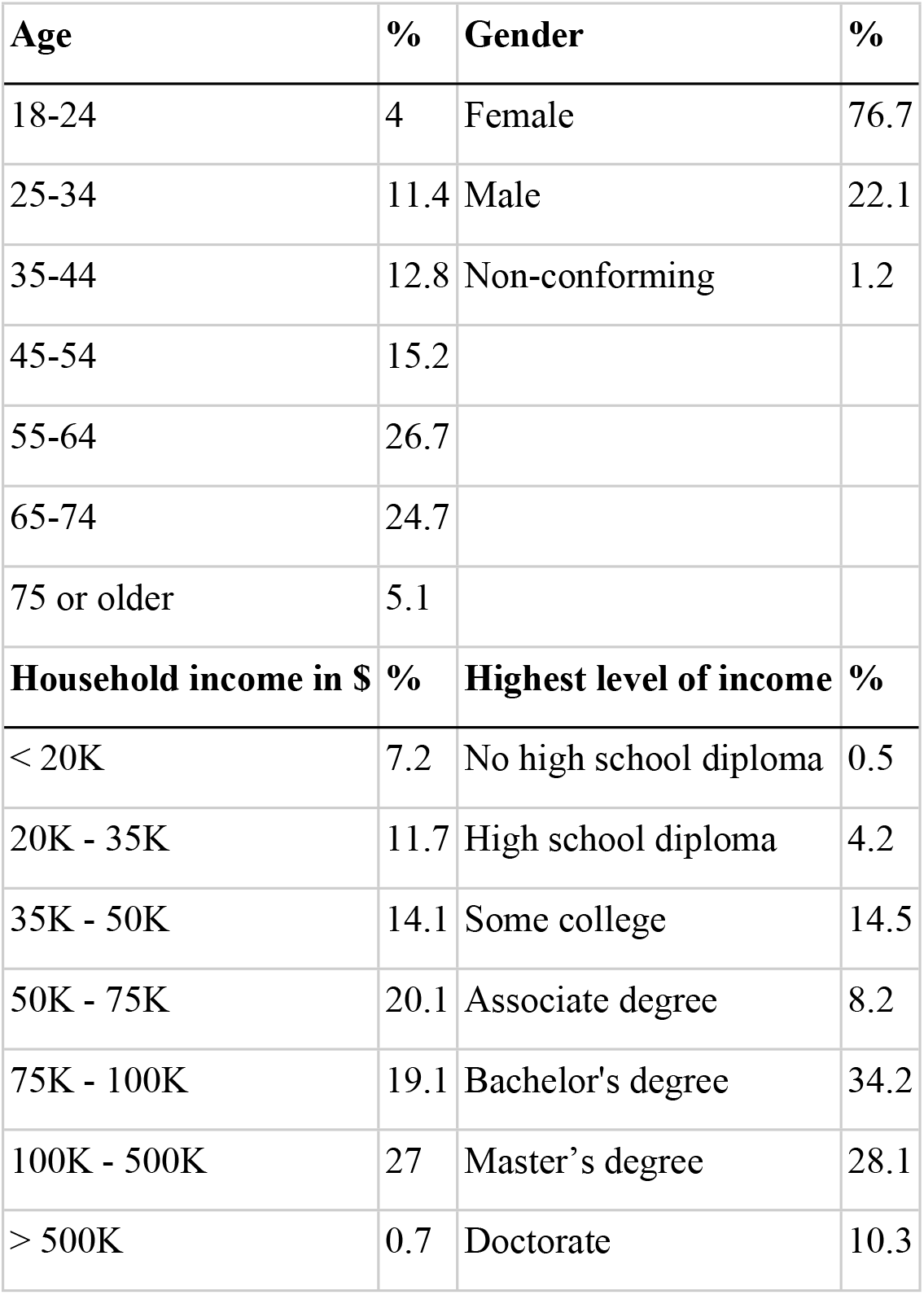
Basic Demographics. Demographic summary of survey respondents.

| Age | % | Gender | % |
| --- | --- | --- | --- |
| 18-24 | 4 | Female | 76.7 |
| 25-34 | 11.4 | Male | 22.1 |
| 35-44 | 12.8 | Non-conforming | 1.2 |
| 45-54 | 15.2 |  |  |
| 55-64 | 26.7 |  |  |
| 65-74 | 24.7 |  |  |
| 75 or older | 5.1 |  |  |
| Household income in \$ | % | Highest level of income | % |
| < 20K | 7.2 | No high school diploma | 0.5 |
| 20K - 35K | 11.7 | High school diploma | 4.2 |
| 35K - 50K | 14.1 | Some college | 14.5 |
| 50K - 75K | 20.1 | Associate degree | 8.2 |
| 75K - 100K | 19.1 | Bachelor's degree | 34.2 |
| 100K - 500K | 27 | Master's degree | 28.1 |
| > 500K | 0.7 | Doctorate | 10.3 |

**Table 2.** Further Demographics. Summary of responses to various questions related to lawns and wildflowers.

| **Owner of a grass lawn** | **%** | **Member of wildflower or pollinator organization** | **%** |
| --- | --- | --- | --- |
| Yes | 82.8 | Yes | 40.9 |
| No | 17.2 | No | 59.1 |
| **Resident of HOA** | **%** | **Already created wildflower plot** | **%** |
| Yes | 21.2 | Yes | 57.1 |
| No | 78.8 | No | 42.9 |
| **Would you create a 6 x 6 ft. wildflower plot** | **%** | **Are you concerned about pollinator declines** | **%** |
| Definitely no | 0.8 | Not at all | 1.1 |
| Probably no | 2.8 | Slightly | 1.0 |
| Unsure | 3.8 | Moderately | 6.7 |
| Probably yes | 14.1 | Very | 20.0 |
| Definitely yes | 78.5 | Extremely | 71.1 |

### What barriers are most important in preventing people from converting patches of lawns to wildflowers?

The most selected answer was that none of the barriers would prevent respondents from converting lawns to wildflowers (36.4% of respondents; Fig. 1). Two of the personal barriers were the highest, ‘Maintenance time’ (27.8%) and ‘Not knowing what to do’ (27.0%). Other personal factors had lower response rates: ‘Maintenance cost’ (12.6%), ‘Appearance’ (8.3%), and ‘Loss of space for recreation’ (3.8%) (Fig. 1). ‘Undesirable plants’ was the third highest response rate (15.1%) but external barriers related to animals were both low: ‘Undesirable wildlife’ (3.4%) and ‘Bee stings’ (2.6%) (Fig. 1). External barriers from other people all had intermediate response rates: ‘Fines or infractions from local government’ (13.9%), ‘Violations of homeowner association policies’ (12.8%) and ‘Opinions of neighbors’ (9.1%).

**Fig. 1.**
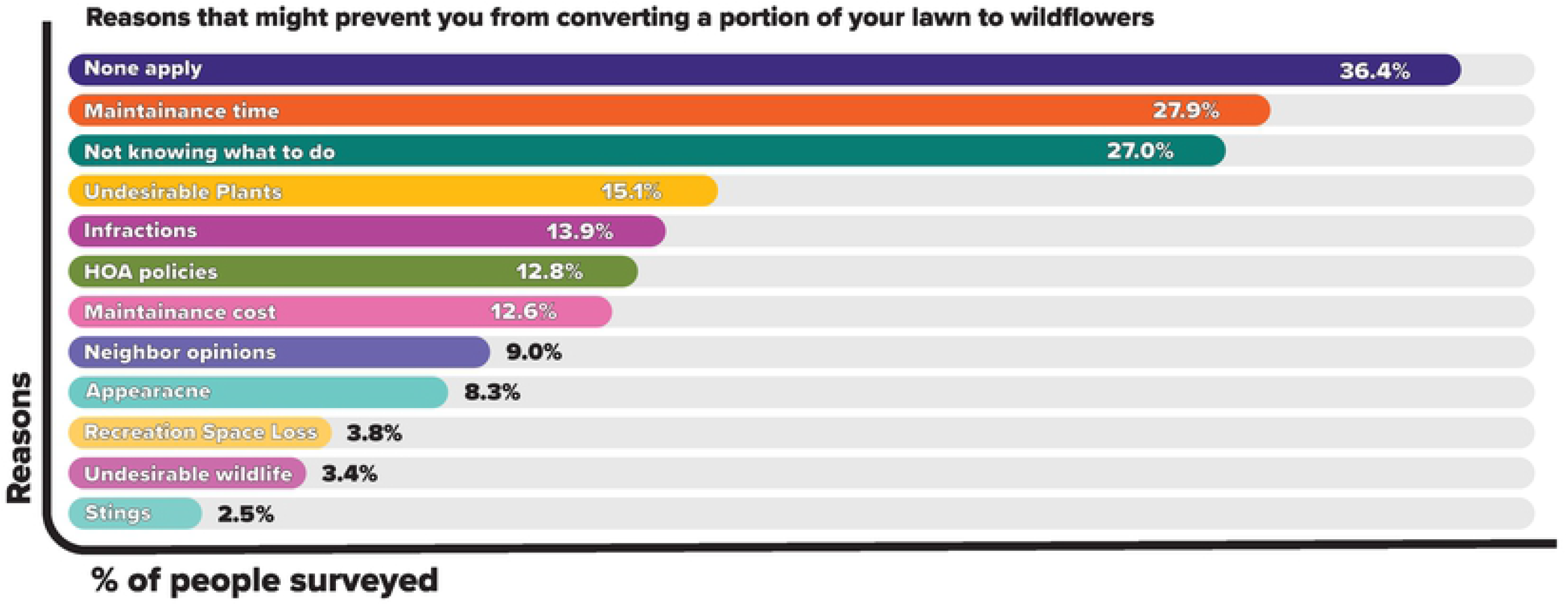
Barriers to Lawn to Wildflower Conversion. Percent of responses to the question: “Of the following items, select those that might prevent you from converting a portion of your lawn to wildflowers (you may select multiple items)”. Items were rearranged to be in descending order based on percentage of responses.

### How do results vary across demographic factors and personal characteristics?

We found that several demographic factors had large impacts on survey responses, especially age and income (Figs. 2 and 3, Table 3). For eight out of 11 barriers age had a highly significant impact, and for *all* barriers we found that younger people were more likely to say at least one barrier would prevent lawn to wildflower conversion than older people (Fig. 2, Table 3). People aged 18-34 were 8.4 times more likely to say that ‘Loss of space for recreation’ was as a barrier than those 65 or older (Fig. 2). This same age group was 4.3 times more likely to indicate ‘Maintenance cost’ (Fig. 2, Table 3), 3.1 times for ‘Fines or infraction from local government’ (Table 3), 3.1 times for ‘Bee stings’ (Table 3), 2.5 times for ‘Violation of HOA policies’ (Table 3), 2.1 times for ‘Undesirable plants’ (Table 3), 2.0 times for ‘Not knowing what to do’ (Fig. 2, Table 3), and 1.7 times more likely to indicate ‘Maintenance time’ as potential barriers (Fig. 2, Table 3). Income also shaped responses, with people living in households making $35K a year or less being 2.2 times more likely to list ‘Maintenance cost’, and 1.6 times most likely to list ‘Fines from local government’ as barriers than people making $75K to $500K per year (Fig. 3, Table 3). Conversely, we found that households making $75K to $500K per year being 2.2 times more likely to select ‘Appearance’ as a barrier than households making less than $35K (Fig. 3, Table 3).

**Fig 2.**
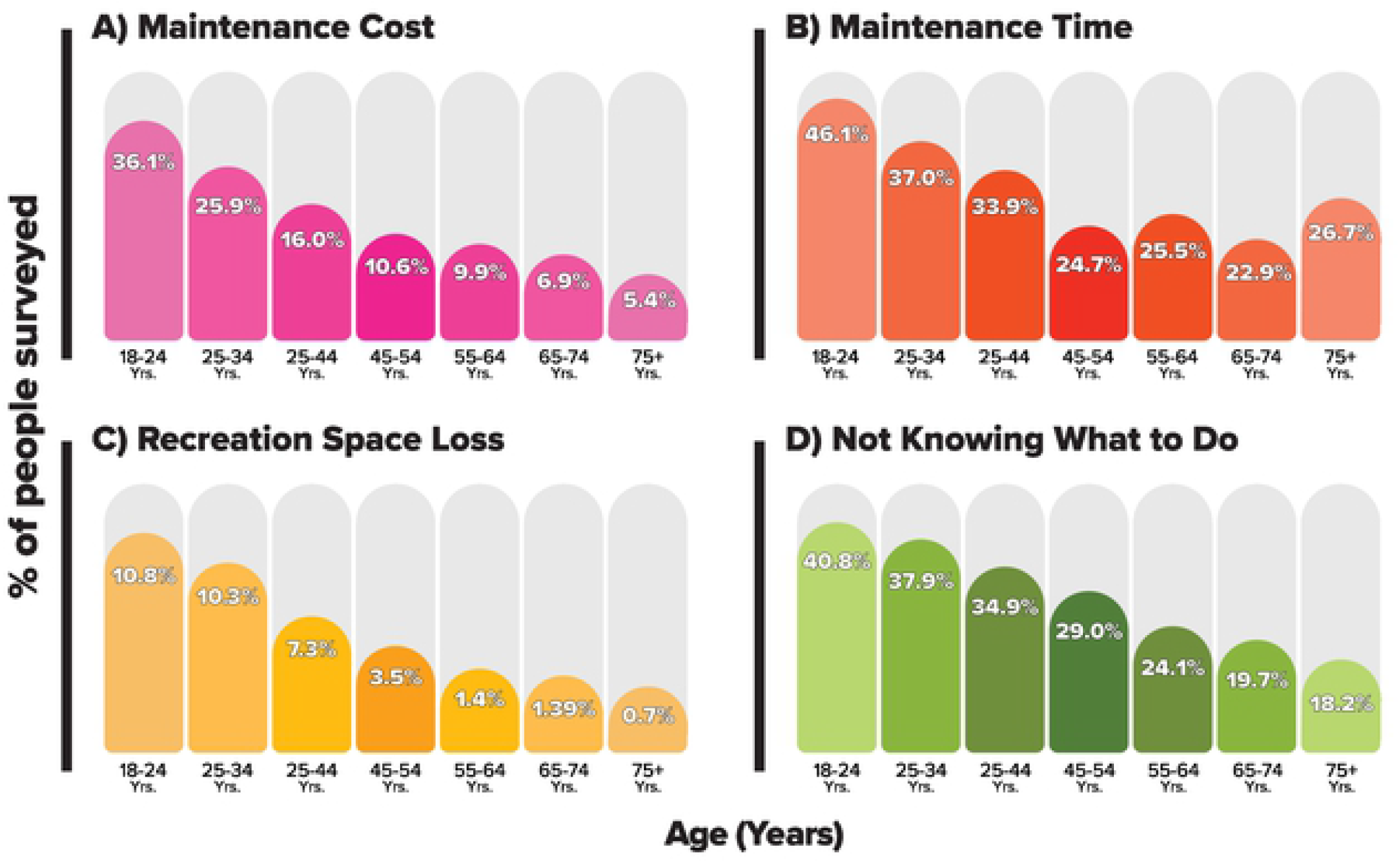
Effects of Age on Survey Results. Percentage of respondents, separated by age, who said that these factors might prevent them from converting a portion of your lawn to wildflowers. The factors shown are A) ‘Maintenance cost’, B) ‘Maintenance time’, C) ‘Loss of space for recreation’, and D) ‘Not knowing what to do’.

**Table 3.** Impacts of Demographics on Survey Responses. Results testing the independence among various demographic factors and the counts of respondents who said each factor might prevent them from converting a portion of their lawn to wildflowers. Table shows χ^2^ test statistic and asterisks or bold font to indicate P-values (*<0.05, **<0.01, and entries with P < 0.001 are bolded).

| Reason | Age | Gender | Income | Education | Member of a HOA | Member of wildflower or pollinator org. | Already made wildflower plot |
| --- | --- | --- | --- | --- | --- | --- | --- |
| Maintenance cost | **166.15** | 0.57 | **48.05** | 5.66 | 1.40 | **41.62** | **68.63** |
| Maintenance time | **59.054** | 0.02 | 4.59 | **26.64** | 9.44** | 9.38** | **58.57** |
| Bee stings | **36.37** | 0.03 | 2.65 | 13.10* | 0.31 | **25.86** | **36.15** |
| Undesirable wildlife | 20.56** | 0.52 | 3.07 | 12.86* | 5.59* | **15.86** | **20.35** |
| Undesirable plants | **47.51** | 0.12 | 12.91* | 4.86 | 0.76 | **17.97** | **40.46** |
| Loss of space for recreation | **104.61** | 0.36 | 7.57 | 3.63 | 0.15 | 6.44* | **22.23** |
| Not knowing what to do | **79.69** | **16.11** | 8.34 | 9.02 | 0.03 | **81.93** | **216.56** |
| Appearance | 13.82* | 0.55 | **28.75** | 9.11 | 7.84** | **15.46** | **23.83** |
| Opinions of neighbors | 21.59** | 0.03 | 12.17* | 4.49 | **81.54** | 0.02 | 1.50 |
| Violation of HOA policies | **59.22** | 0.45 | 1.30 | 8.46 | **760.65** | 1.11 | 5.58* |
| Infractions from local government | **95.12** | 0.94 | **22.30** | 6.18 | 6.57* | 5.72* | 4.11* |
| None of these apply to me | **221.14** | 7.79** | 7.87 | 10.85 | **53.42** | **62.38** | **162.08** |

Other demographics had smaller effects on responses, such as gender, education level, and membership in an HOA. We found that women were 1.4 times more likely to say that ‘Not knowing what to do’ would prevent them from participating (Fig. 4, Table 3). This was the only barrier that gender played a significant role in the response. People with higher education levels selected ‘Maintenance time’ more than those with less education (Fig. 4, Table 3). Specifically, those with a doctorate degree were 1.9 times more likely to select ‘Maintenance time’ than those with high school education (Fig. 4). Finally, people in homeowners’ associations were 11.5 times more likely to select ‘Violation of HOA policies’ and 2.8 more likely to select ‘Opinions of neighbors’ as potential barriers (Table 3, Fig. 4).

**Fig 3.**
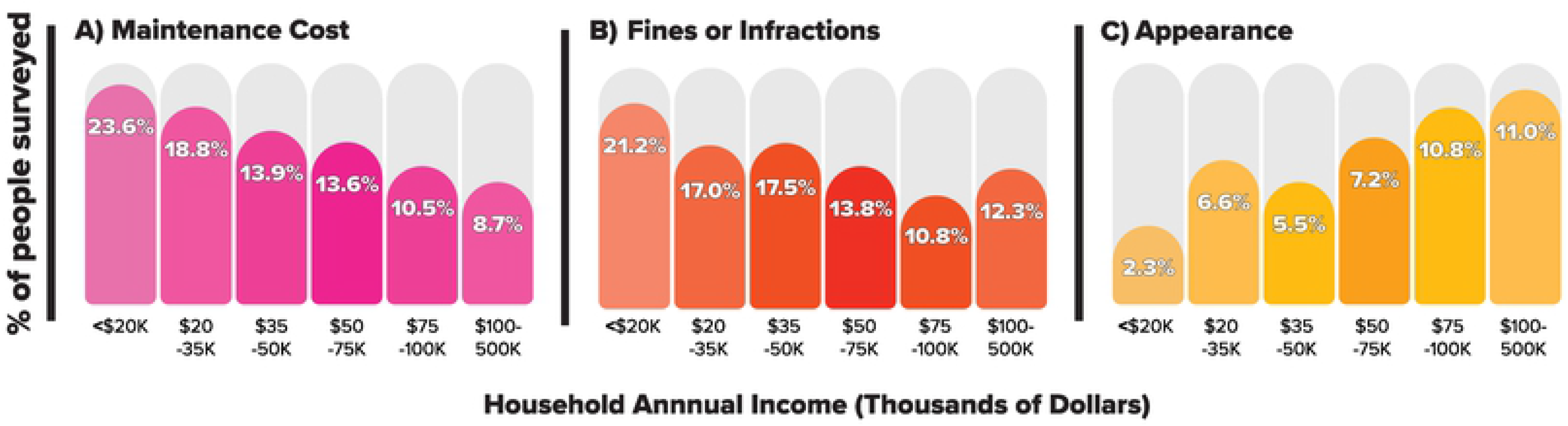
Effects of Income on Survey Results. Percentage of respondents, separated by household income, who said that these factors might prevent them from converting a portion of your lawn to wildflowers. The factors shown are A) ‘Maintenance cost’, B) ‘Appearance’, and C) ‘Fines or infractions from local government’.

**Fig 4.**
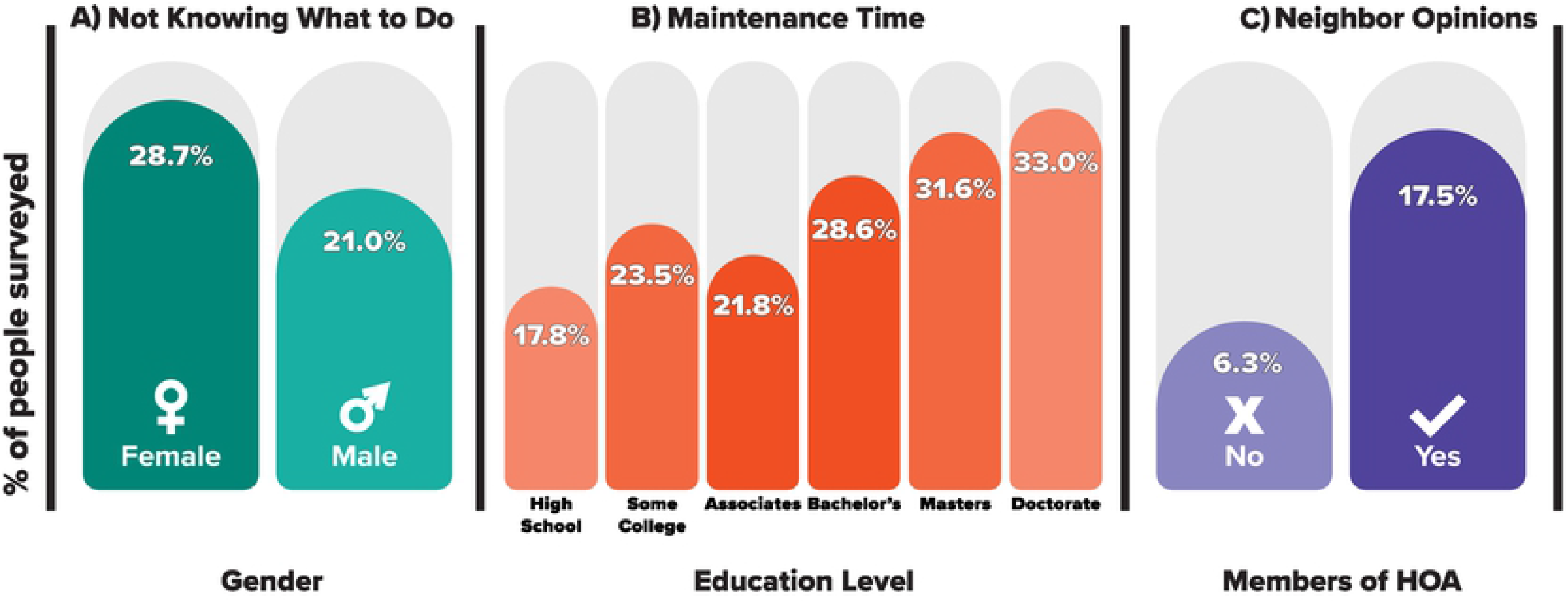
Other Demographic Factors on Survey Results. Percentage of respondents who said that these barriers might prevent them from converting a portion of your lawn to wildflowers. The barriers shown are A) ‘Not knowing what to do’ and how responses vary across gender, B) ‘Maintenance time’ and how responses vary across income level, and C) ‘Opinions of neighbors’ and how responses vary across membership to a homeowner’s association.

Respondents who were either a member of a native plant or pollinator organization or had already created a wildflower plot selected fewer barriers overall than other respondents (Tables 2 and 3). Membership in native plant or pollinator organization reduced number of barriers selected for six of 11 potential barriers (Table 3). And having previously created a plot strongly reduced number of barriers selected for eight out of 11 potential barriers (Table 3). Specifically, membership in a native plant or pollinator organization reduced likelihood of selecting ‘Not knowing what to do’ by 1.7 times and those who had already created a wildflower plot reduced likelihood of this response by 2.7 times (Table 3). The only three barriers that were not impacted by if the respondent had already made a wildflower plot were external factors from other people: ‘Opinions of neighbors’, ‘Violation of HOA policies’, and ‘Fines or infractions from local government’ (Table 3).

## Discussion

We surveyed over 3200 people about what factors might prevent them from converting grass lawns to native wildflowers. Survey respondents were from around the United States and Canada but did not represent a random sample of the general public. Instead they represented members of native plant societies, other plant and pollinator organizations, or people who were interested in plants and concerned about pollinator declines (see Table 2). This audience skewed heavily female (77%), and was older, more educated, and higher income than the general population (Table 1, S1 Table). Among the 11 barriers we included that could prevent lawn to wildflower conversions, most (9 of 11) had fewer than 15% of people select them. The most common response in the survey was “None apply to me” (36.4% of people). This suggests that the population we surveyed was, overall, keen to convert lawns to wildflowers. In fact, 57% of respondents had already created wildflower plots (Table 2) and our results should be interpreted accordingly. By taking that into consideration, our survey results point to two main conclusions: 1) ‘Maintenance time’ and ‘Not knowing what to do’ are the most important barriers to lawn to wildflower conversions, and 2) age and income play large roles in shaping barriers to creating wildflower plots.

### Most important barriers to creating wildflower plots

The two most common barriers to homeowners creating a wildflower plot were ‘Maintenance time’ and ‘Not knowing what to do’. ‘Maintenance time’ has previously been found to be a significant concern of homeowners when surveyed about lawn management decisions [18, 22, 25] as respondents value a landscape design that allows them to enjoy their yard with minimal impact on their already limited time. ‘Not knowing what to do’ may have been an important barrier for two reasons. First, lawns and landscapes featuring native plants are not common [26] and often not an accepted part of community culture [20, 27]. Therefore, knowledge about creating or maintaining such landscapes may not be commonplace. However, our results are skewed by a larger percentage of respondents involved in plant and pollinator organization, so it is likely that the general population would have chosen ‘Not knowing what to do’ more frequently. Our data supports this idea as 33% of our respondents who were not part of a native plant or pollinator organization selected ‘Not knowing what to do’ as a barrier compared to 18% for members. Second, there are many different methods for converting lawns to native plants [28] and numerous native plants to choose from that require different growing conditions. This excess of information and options could lead to confusion or analysis paralysis [29]. Outreach efforts by Xerces Society (https://xerces.org/) and Lawn to Wildflowers (https://lawntowildflowers.org) are attempting to address these two barriers by presenting simple and clear protocols for creating wildflower habitat, and resources for selecting plants and seeds that are not overwhelming.

Another significant barrier to plot creation related to the potential opinions and objections of the homeowner’s local governments, HOAs, and neighbors. These concerns, while not as common as those regarding ‘Maintenance time’ and ‘Not knowing what to do’, persisted even within those individuals who already creased a wildflower plot. Previous studies into the cultural norms surrounding US yards have found that the types of yards that neighbors had significantly affect how a homeowner designs their own yard [19]. Finding a method that allows homeowners to incorporate ecologically beneficial features such as wildflower plots in a way that does not compromise the propriety of their neighbors will be essential [23]. An important thing to note is we expect the general public’s concerns about external factors (such as neighbors) to be stronger than what we found, which could be difficult obstacle for programs promoting pollinator-friendly lawns. This could be because our surveyed audience was presumably more open to these concepts, as they largely belonged to native plant societies and have some background knowledge in the importance of urban ecology. Further examination of the perception of those outside of our skewed audience will be essential when engaging the public with initiatives encouraging the creation of wildflower plots within traditional lawns [30].

‘Undesirable plants’ was the most selected nature-related barrier and this likely reflects the topic-specific knowledge and experience of our audience since weeds are a serious problem in plant restoration. Restoration experts say that non-native or invasive plants are the single biggest threat to success when restoring native prairies [31]. Therefore, it is actually quite encouraging that only 15% of people say that might prevent them from creating a wildflower plot since it is such threat to success. Still, it does suggest that promoting methods that could help suppress weeds could be an important tactic for outreach. Other nature barriers such as ‘Undesirable wildlife’ and ‘Bee stings’, as well as ‘Loss of space for recreation’, could also be more important to the general public, but our results suggest that these are of minimal concern to our audience and probably not something that native plant and pollinator organizations need to address.

### Impacts of age and income on barriers to lawn to wildflower restorations

Age was the most important demographic factor shaping our results. Across all barriers, younger people were more likely to be dissuaded than older people, the most dramatic example being with ‘Maintenance cost’ (Fig. 3, Table 3). The reasons for these patterns are not clear but there are a few possible reasons. First, some argue that younger generations on average spend less time outside and may be less interested and concerned about nature and the environment [32]. If younger people are less interested in nature and native plants that could make them more easily dissuaded from creating a wildflower plot. Second, younger generations have lower incomes, are less likely to own homes, and may be more likely to move [33]. It makes sense that all these factors would make younger people less able to actually create wildflower gardens, and perhaps these life experiences, also make them more pessimistic when imagining what would prevent them given the hypothetical situations we asked about. This highlights two major difficulties in reaching out to younger people: they likely have much less opportunities and resources to plant native wildflowers, and presumably are more easily discouraged from doing so even if they had the means. However, our results suggest that promoting lawn to wildflower methods that are clear, simple, and inexpensive could be helpful.

Income, education, gender, and membership in HOA’s and plant and pollinators organizations also shaped responses. The results we found related to income were predictable and understandable. People with less money are more likely to be concerned with cost and fines from local governments. Interestingly, those with higher incomes were more concerned with appearances (Fig. 3). This could be because high income people live in neighborhoods where well or professionally manicured lawns are commonplace, and that may shape views on beauty expectations for yards [19]. Men were less deterred by ‘Not knowing what to do’ then women, but the reasons for this small effect are not known. Not surprisingly, people in HOAs were 11.5 times more likely to be deterred by HOA policies, but they were also 2.8 times more likely to be deterred by the ‘Opinions of neighbors’. These results reinforce the idea that the influence of HOAs result in communities that self-enforce strong social norms regarding appearances of lawns [34].

## Conclusions

One of the primary motivations for this study was to guide the outreach efforts of our public science project Lawn to Wildflowers and other organizations that are advocating for native plants and pollinators. Given the results of our survey we have the following recommendations for organizations promoting pollinator-friendly native plant gardens:

- Promote easy-to-maintain landscapes, and make clear that native plant landscapes could result in less maintenance time than mowed turfgrass.
- Give clear instructions on creating wildflower plots with only a few options. Instructions should be specific, easy to follow, and do not require purchasing specialized equipment.
- Provide, or link to, native plants guides or seed sources that have few enough options to not be overwhelming.
- Target older audiences.
- When targeting a younger audience, focus on promoting small wildflower plots that are cheaper to create, less time consuming to maintain, and simpler to give easy and specific instructions on creating. Alternative approaches using moveable pots or containers may also be more accessible and appealing.
- For lower income audiences focus on more cost-effective approaches like sowing seeds, and for higher income audience suggest more expensive options of larger potted plants to transplant, which may also have more attractive appearances.

## Acknowledgements

We would like to that Bobby Jeanpierre and for assistance with writing the survey, Omar Abdellatif for creating the figures, Bree Goldstein for guidance on online marketing, and Shiala Morales for assistance throughout the project. We are grateful for funding for this project, provided by the Foundation for Food and Agriculture, Pollinator Health Fund (Grant ID: 549058) to BJS and NET.

## Supporting information

- S1 Appendix. Complete text of the online survey.
- S2 Appendix. The full dataset used in the final analyses.
- S1 Figure. Map showing location of 3249 people who took our online survey.
- S2 Figure. Advertisement that ran on Facebook and Instagram to promote our online survey.
- S1 Table. United States Census data from 2018 to compare to the demographic data collected in our online survey.

## References

1. Foley JA, DeFries R, Asner GP, Barford C, Bonan G, Carpenter SR, et al. Global consequences of land use. Science. 2005;309(5734):570–4. doi: 10.1126/science.1111772.

2. Newbold T, Hudson LN, Hill SL, Contu S, Lysenko I, Senior RA, et al. Global effects of land use on local terrestrial biodiversity. Nature. 2015;520(7545):45–50. doi: 10.1038/nature14324.

3. Rosenberg KV, Dokter AM, Blancher PJ, Sauer JR, Smith AC, Smith PA, et al. Decline of the North American avifauna. Science. 2019;366(6461):120–4. doi: 10.1126/science.aaw1313.

4. Goulson D, Nicholls E, Botías C, Rotheray EL. Bee declines driven by combined stress from parasites, pesticides, and lack of flowers. Science. 2015;347(6229):1255957. doi: 10.1126/science.1255957.

5. Koh I, Lonsdorf EV, Williams NM, Brittain C, Isaacs R, Gibbs J, et al. Modeling the status, trends, and impacts of wild bee abundance in the United States. Proc Natl Acad Sci U S A. 2016;113(1):140–5. doi: 10.1073/pnas.1517685113.

6. Potts SG, Biesmeijer JC, Kremen C, Neumann P, Schweiger O, Kunin WE. Global pollinator declines: trends, impacts and drivers. Trends Ecol Evol. 2010;25(6):345–53. doi: 10.1016/j.tree.2010.01.007.

7. Sánchez-Bayo F, Wyckhuys KA. Worldwide decline of the entomofauna: A review of its drivers. Biol Conserv. 2019;232:8–27. doi: 10.1016/j.biocon.2019.01.020.

8. Gallai N, Salles J-M, Settele J, Vaissière BE. Economic valuation of the vulnerability of world agriculture confronted with pollinator decline. Ecol Econ. 2009;68(3):810–21. doi: 10.1016/j.ecolecon.2008.06.014.

9. Kearns CA, Inouye DW, Waser NM. Endangered mutualisms: the conservation of plant-pollinator interactions. Annu Rev Ecol Syst. 1998;29(1):83–112. doi: 10.1146/annurev.ecolsys.29.1.83.

10. Kremen C, Williams NM, Thorp RW. Crop pollination from native bees at risk from agricultural intensification. Proc Natl Acad Sci U S A. 2002;99(26):16812–6. doi: 10.1073/pnas.262413599.

11. Smith LS, Broyles MEJ, Larzleer HK, Fellowes MDE. Adding ecological value to the urban lawnscape. Insect abundance and diversity in grass-free lawns. Biol Conserv. 2015;24(1):47–62. doi: 10.1007/s10531-014-0788-1.

12. Cristina M, Running S, Elvidge C, Dietz J, Tuttle B, Nemani R. Mapping and modeling the biogeochemical cycling of turf grasses in the United States. Environ Manage. 2005;36:426–38. doi: 10.1007/s00267-004-0316-2.

13. Milesi C, Elvidge CD, Nemani RR. Assessing the extent of urban irrigated areas in the United States. Boca Raton, Fla.: CRS Press; 2009.

14. USDA. National Agricultural Statistics Service, 2012 Census, Volume 1, Chapter 1: U.S. National Level Data. Table 36; 2012. Available from: https://www.nass.usda.gov/Publications/AgCensus/2012/Full_Report/Volume_1,_Chapter_1_US/st99_1_034_036.pdf.

15. Goddard MA, Dougill AJ, Benton TG. Scaling up from gardens: biodiversity conservation in urban environments. Trends Ecol Evol. 2010;25(2):90–8. doi: 10.1016/j.tree.2009.07.016.

16. Larson JL, Dale A, Held D, McGraw B, Richmond DS, Wickings K, et al. Optimizing pest management practices to conserve pollinators in turf landscapes: current practices and future research needs. J Integr Pest Manag. 2017;8(1):18. doi: 10.1093/jipm/pmx012.

17. Hugie K, Yue C, Watkins E. Consumer preferences for low-input turfgrasses: A conjoint analysis. HortScience. 2012;47(8):1096–101. doi: 10.21273/HORTSCI.47.8.1096.

18. Larson KL, Nelson KC, Samples SR, Hall SJ, Bettez N, Cavender-Bares J, et al. Ecosystem services in managing residential landscapes: priorities, value dimensions, and cross-regional patterns. Urban Ecosyst. 2016;19(1):95–113. doi: 10.1007/s11252-015-0477-1.

19. Nassauer JI, Wang Z, Dayrell E. What will the neighbors think? Cultural norms and ecological design. Landsc Urban Plan. 2009;92(3-4):282–92. doi: 10.1016/j.landurbplan.2009.05.010.

20. Zheng B, Zhang Y, Chen J. Preference to home landscape: wildness or neatness? Landsc Urban Plan. 2011;99(1):1–8. doi: 10.1016/j.landurbplan.2010.08.006.

21. Beard JB, Green RL. The role of turfgrasses in environmental protection and their benefits to humans. J Environ Qual. 1994;23(3):452–60. doi: 10.2134/jeq1994.00472425002300030007x.

22. Larson KL, Casagrande D, Harlan SL, Yabiku ST. Residents’ yard choices and rationales in a desert city: social priorities, ecological impacts, and decision tradeoffs. Environ Manage. 2009;44(5):921. doi: 10.1007/s00267-009-9353-1.

23. Nassauer JI. Messy ecosystems, orderly frames. Landsc J. 1995;14(2):161–70. doi: 10.3368/lj.14.2.161.

24. Dahmus ME, Nelson KC. Yard stories: examining residents’ conceptions of their yards as part of the urban ecosystem in Minnesota. 2014;17(1):173–94. doi: 10.1007/s11252-013-0306-3.

25. Martini NF, Nelson KC, Hobbie SE, Baker LA. Why “feed the lawn”? Exploring the influences on residential turf grass fertilization in the Minneapolis-Saint Paul metropolitan area. Environ Behav. 2015;47(2):158–83. doi: 10.1177/0013916513492418.

26. Helfand GE, Park JS, Nassauer JI, Kosek S. The economics of native plants in residential landscape designs. Landsc Urban Plan. 2006;78(3):229–40. doi: 10.1016/j.landurbplan.2005.08.001.

27. Gobster PH, Nassauer JI, Daniel TC, Fry G. The shared landscape: what does aesthetics have to do with ecology? Landsc Ecol. 2007;22(7):959–72. doi: 10.1007/s10980-007-9110-x.

28. Sarah Foltz Jordan, Jessa Kay Cruz, Kelly Gill, Jennifer Hopwood, Jarrod Fowler, Eric Lee-Mäder, et al. Organic Site Preparation for Wildflower Establishment. 2016. Available from: https://xerces.org/publications/guidelines/organic-site-preparation-for-wildflower-establishment.

29. Iyengar SS, Lepper MR. When choice is demotivating: Can one desire too much of a good thing? J Pers Soc Psychol. 2000;79(6):995. doi: 10.1037/0022-3514.79.6.995.

30. Groffman PM, Stylinski C, Nisbet MC, Duarte CM, Jordan R, Burgin A, et al. Restarting the conversation: challenges at the interface between ecology and society. Front Ecol Environ. 2010;8(6):284–91. doi: 10.1890/090160.

31. Rowe HI. Tricks of the trade: techniques and opinions from 38 experts in tallgrass prairie restoration. Restor Ecol. 2010;18:253–62. doi: 10.1111/j.1526-100X.2010.00663.x.

32. Louv R. Last child in the woods: Saving our children from nature-deficit disorder: Algonquin books; 2008.

33. McKee K. Young people, homeownership and future welfare. Hous Stud. 2012;27(6):853–62. doi: 10.1080/02673037.2012.714463.

34. Fraser JC, Bazuin JT, Band LE, Grove JM. Covenants, cohesion, and community: The effects of neighborhood governance on lawn fertilization. Landsc Urban Plan. 2013;115:30–8. doi: 10.1016/j.landurbplan.2013.02.013.

